# Sex-biased response to and brain cell infection by SARS-CoV-2 in a highly susceptible human ACE2 transgenic model

**DOI:** 10.1101/2021.05.04.441029

**Authors:** Ching-Yen Tsai, Chiung-Ya Chen, Jia-Tsrong Jan, Yu-Chi Chou, Mei-Ling Chang, Lu A Lu, Pau-Yi Huang, Mandy F.-C. Chu, Tsan-Ting Hsu, Yi-Ping Hsueh

**Author notes:** Co-corresponding authors: Ching-Yen Tsai, and Yi-Ping Hsueh.

## Abstract

The COVID-19 pandemic is caused by SARS-CoV-2 infection. Human angiotensin-converting enzyme II (hACE2) has been identified as the receptor enabling SARS-CoV-2 host entry. To establish a mouse model for COVID-19, we generated transgenic mouse lines using the (HS4)_2_-pCAG-hACE2-HA-(HS4)_2_ transgene cassette, which expresses HA-tagged hACE2 under control of the CAG promoter and is flanked by HS4 insulators. Expression levels of the hACE2 transgene are respectively higher in lung, brain and kidney of our CAG-hACE2 transgenic mice and relatively lower in duodenum, heart and liver. The CAG-hACE2 mice are highly susceptibility to SARS-CoV-2 infection, with 100 PFU of SARS-CoV-2 being sufficient to induce 87.5% mortality at 9 days post-infection and resulting in a sole (female) survivor. Mortality was 100% at the higher titer of 1000 PFU. At lower viral titers, we also found that female mice exposed to SARS-CoV-2 infection suffered much less weight loss than male mice, implying sex-biased responses to SARS-CoV-2 infection. We subjected neuronal cultures to SARS-CoV-2 pseudovirus infection to ascertain the susceptibilities of neurons and astrocytes. Moreover, we observed that expression of SARS-CoV-2 Spike protein alters the synaptic responses of cultured neurons. Our transgenic mice may serve as a model for severe COVID-19 and sex-biased responses to SARS-CoV-2 infection, aiding in the development of vaccines and therapeutic treatments for this disease.

## Introduction

Since the outbreak of Coronavirus disease 2019 (COVID-19), more than 84 million cases have been confirmed worldwide and almost 1.8 million people have died (https://covid19.who.int/, WHO Coronavirus Disease (COVID-19) Dashboard, 2021/1/4 updated). COVID-19 is a novel infectious disease caused by SARS-CoV-2 virus [1, 2]. SARS-CoV-2 binds to the receptors angiotensin-converting enzyme 2 (ACE2) and neuropilin-1 (NRP1) on host cells via its Spike envelope glycoprotein [3]. Small animal models that respond to SARS-CoV-2 infection and mimic resulting clinical symptoms and pathologies are crucial to model COVID-19 for mechanistic study, vaccine development and potential therapy.

To date, mustelids, some felids and rodents, and various nonhuman primates have exhibited susceptibility to SARS-CoV-2 infection. However, infection in those species does not result in clinical diseases resembling those reported for human [4, 5]. Mice are not susceptible to SARS-CoV-2 infection because of *Ace2* sequence variation [5] Thus, establishing a genetically-modified mouse model that expresses hACE2 would provide valuable insights into clinically-relevant disease pathogenesis. Accordingly, several mouse models have already been established to study COVID-19. First, K18-ACE2 transgenic mice that express a hACE2 transgene under the control of the human K18 promoter were established for SARS-CoV study a decade ago, representing the first available mouse model for SARS-CoV-2 infection [6, 7]. Later, adenovirus type 5 was used to deliver and transiently express hACE2 in mice [8, 9], and inducible hACE2 expression was established in rtTA-hACE2 transgenic mice [10] for SARS-CoV-2 study. HFH4-hACE2 transgenic mice that express a hACE2 transgene driven by the hepatocyte nuclear factor-3/forkhead homologue 4 (HFH4) promoter have since been developed [11]. In addition to ectopic expression of hACE2 under the control of other gene promoters, mAce2-hACE2 transgenic mice [12] and hACE2-knockin mice [13] in which the mouse *Ace2* promoter is used to drive hACE2 transgene expression have also been generated.

Studies using the aforementioned animal models have established systems for investigating SARS-CoV-2 infection in a variety of tissues and explored the immune response to viral challenge. However, the sex-biased responses observed among human patients have not been investigated in mouse models. In this study, we first established a transgenic mouse line in which hACE2 is ubiquitously expressed under the control of the CAG promoter. Our transgenic mice proved highly susceptible to SARS-CoV-2 challenge, with a virus titer of 100 PFU being sufficient to induce lethality. Moreover, our female and male transgenic mice exhibited differential responses to SARS-CoV-2 infection. Finally, we also show that cultured neurons and glial cells prepared from our CAG-hACE2 transgenic mice were readily infected by pseudovirus expressing SARS-CoV-2 Spike proteins. In conclusion, this new mouse line serves as an appropriate model for severe COVID-19 disease, enabling investigations of sex-biased responses to SARS-CoV-2 infection and its impact on various tissues, including the brain. Our transgenic model will be valuable for vaccine development and evaluating COVID-19 therapies.

## Materials and Methods

### Animals

Mice were bred and maintained in the animal facility of the Institute of Molecular Biology (IMB), Academia Sinica, under pathogen-free conditions. The mice were group-housed with their littermates and each cage contained 3 to 5 mice. All animal experiments were performed with the approval of the Academia Sinica Institutional Animal Care and Utilization Committee (IACUC Protocol No. 12-08-391) and in strict accordance with its guidelines and those of the Council of Agriculture Guidebook for the Care and Use of Laboratory Animals. Viral challenge experiments with SARS-CoV-2 were performed in the P3 animal facility of the Genomic Research Center. The protocol for animal experiments at the P3 level was also evaluated and approved by the IACUC of Academia Sinica (Protocol No. 20-11-1538). Surviving mice after viral challenge were euthanized using carbon dioxide.

### Plasmid constructions

#### For pseudovirus

To generate pcDNA3.1-nCoV-SΔ18, the SARS-CoV-2 *Spike* gene with a 54-nucleotide deletion at the C-terminus was PCR-amplified from synthetic DNA (provided by Alex Ma, Academia Sinica, Taiwan) using the Kapa HiFi PCR kit (Kapa Biosystems) with a primer pair (Forward, 5’-AACTTAAGCTTGGTACCGCCACCATGTTCGTCTTCCTGGTCCTG-3’; Reverse, 5’-TGCTGGATATCTGCAGAATTCTTACTTACAGCAGGAGCCACAGCTACAGCAG-3’), and subcloned into KpnI and EcoRI sites of pcDNA3.1™(+) expression vector using a GenBuilder™ Cloning kit (GeneScript^®^).

#### For hACE2 transgenic mice

The full-length *hAce2* gene was PCR-amplified from Mammalian Gene Collection cDNA clone (clone number MGC:47598) using a Kapa HiFi PCR kit (Kapa Biosystems) with a primer pair (Forward, 5’-GGGAGACCCAAGCTGGCTAGCCACCATGTCAAGCTCTTCCTGGCTCCTTC-3’ and Reverse, 5’-TTGTCTCAAGATCTAGAATTCCTAAAAGGAGGTCTGAACATCATCAGTG-3’), and subcloned into the NheI and EcoRI sites of pLAS2w.Pbsd (a lentiviral transfer vector from the RNAi Core of Academia Sinica, Taiwan) using a GenBuilderTM Cloning kit (GeneScript^®^). The hACE2 cDNA was further PCR-amplified and sub-cloned into pcDNA3.1 using the following primers: Forward, 5’-GCCCTCTAGGCCACCATGTCAAGCTCTTCCTGG-3’; Reverse, 5’-CTAAGCGGGCGCCACCTGGGAGGTCTCGGTACCAAAGGAGGTCTGAACATCATCAGTGT-3’. HA tag sequences were added to the 3’ end of the hACE2 transgenic cDNA using a Q5^®^ Site-Directed Mutagenesis Kit (NEB#E0554S) according to the manufacturer’s instructions. Primers for site-directed mutagenesis were: Forward, 5’-GTTCCAGATTACGCTTAAGCTTGGATCCGCGTTAAGTTTAAACCGCTG-3’; Reverse, 5’-ATCGTATGGGTATCCAGCGGGCGCCACCTGGGA-3’. The pCAG-hACE2-HA DNA fragment was then subcloned into an insulator (HS4)-containing plasmid, as described previously [14]. To generate transgenic mice, the entire (HS4)_2_-pCAG-hACE2-HA-(HS4)_2_ cassette was digested with NotI and isolated for pronuclei microinjection.

#### For transient expression

The previously described construct GW1-SARS2 Spike-cHA [15] was used for transient expression of full-length SARS-CoV-2 Spike protein into cultured cells.

### Generation of hACE2 transgenic mice

For mice production, we super-ovulated 3-4 week-old C57BL/6J female mice with 3.75-5 i.u. of pregnant mare serum gonadotropin (PMSG, Sigma-Aldrich G4877), followed 46-h later by 3.75-5 i.u. of human chorionic gonadotropin (hCG, Sigma-Aldrich CG1063). Super-ovulated female mice were mated to male mice and one-cell-stage zygotes were collected the following day. The (HS4)_2_-pCAG-hACE2-HA-(HS4)_2_ cassette (2 ng/μl) was microinjected into pronuclei of the zygotes. Injected zygotes were transferred into the oviduct of 0.5-dpc (days post-coitum) pseudo-pregnant ICR female mice. Genotyping was performed using genomic PCR with primers specific for ACE2 cDNA (Forward 5’-GAGACTATGAAGTAAATGGGGTAGATGGC-3’; Reverse 5’-CTTCATTAGCTCCATTTCTTAGCAGAAAAGG-3’). Mice genotyped as having a 582-bp PCR product were identified as hACE2 transgenic mice.

### Copy number analysis of the transgene using droplet digital PCR (ddPCR)

To determine hACE2 transgene copy number, genomic DNA was extracted from mouse tail snips using a QuickGene DNA tissue kit S (KUTABO, Cat no.: DT-S). Purity and concentration of DNA samples were measured using a Nanodrop spectrophotometer (Thermo Fisher Scientific, ND-1000). Genomic DNA was digested with SpeI-HF^®^ (NEB, # R3133S) to completely release individual hACE2 transgene cassettes from possible tandem repeats and/or multiple integration sites in a final volume of 20 μL at 37 °C for 1 h, followed by heat inactivation at 65 °C for 20 mins. *Tbp* (TATA box binding protein, gene ID 21374) was used as the two-copy reference gene. The reaction mixtures contained ddPCR™ Supermix for Probes (no dUTPs; Bio-Rad Laboratories, CA, USA), primers (PrimeTime mini qPCR Assay, 100 rxn, IDT), and template DNA (40 ng) in a final volume of 20 μl. Primers and probes are as follows: *Tbp* Forward: 5’-TTGCTACTGCCTGCTGTT-3’; Reverse: 5’-GGACTTACTCCACAGCCTATTC-3’; *Tbp* 5’ Sun probe: 5’-TTGCTGCTGCTGTCTTTGTTGCTC-3’; *hACE2* transgene Forward: 5’-CTAACGGACCCAGGAAATGTTCAGA-3’; Reverse: 5’-GGTTGTGCAGCATATGCCATATCATAC-3’; *hACE2* 5’ FAM probe: 5’-AAGGGCGACTTCAGGATCCTTATGTGCAC-3’. Each reaction was then loaded into a sample well of an eight-well disposable cartridge (DG8™; Bio-Rad Laboratories) along with 70 μl of droplet generation oil (Bio-Rad Laboratories). Droplets were formed using a QX200™ Droplet Generator (Bio-Rad Laboratories) according to the manufacturer’s instructions. Droplets were then transferred to a 96-well PCR plate, heat-sealed with foil and amplified to the end point using a conventional thermal cycler (95 °C for 5 mins, followed by 40 cycles of 94 °C for 30 s and 58 °C for 1 min, and a final extension at 98 °C for 10 mins). The resulting products were scanned on a QX200 Droplet Reader (Bio-Rad Laboratories), and the data was analyzed using QuantaSoft™ software (Bio-Rad Laboratories).

### SARS-CoV-2 infection

CAG-hACE2 transgenic or wild-type (WT) mice were anesthetized and intranasally challenged with SARS-CoV-2 TCDC#4 (hCoV-19/Taiwan/4/2020 obtained from Taiwan Centers of Disease Control; lot: IBMS20200819) in a volume of 100 μL of sterile PBS at the indicated plaque-forming units (PFU).

### Western blot analysis

Transgenic and WT mice were sacrificed before collecting different tissues for lysis. Western blot analysis was performed as described previously [14]. Antibodies and respective dilutions are as follows: hACE2 (Abcam ab108209, 1:2500); HA (Cell Signaling #3725, 1:1000); and HSP90 (provided by Dr. Chung Wang, 1:5000) [16].

### Immunohistochemistry

Transgenic and WT mice were sacrificed before collecting different tissues for fixation in 10% formaldehyde (MACRON H121-08) at 4 °C for more than 24 h. Fixed tissues were stored in 70% ethanol before embedding in paraffin. Tissues were further processed using a Leica TP1020 Semi-enclosed Benchtop Tissue Processor.

Immunostaining was performed as described previously [17]. The following commercial antibodies were used as primary antibodies for immunostaining: rabbit anti-ACE2 (Abcam, ab108209); mouse anti-HA (Abcam, ab130275); and rabbit anti-HA (Cell Signaling, 3742). Tissue sections were colored via DAB-staining (3, 3-diaminobenzidine; Dako, K3468).

### Production of pseudotyped SARS-CoV-2 lentivirus

The pseudotyped SARS-CoV-2 lentivirus, which carries SARS-CoV-2 Spike protein as viral envelope protein, was generated by transiently transfecting HEK-293T cells with pCMV-ΔR8.91, pLAS2w.EGFP.Puro and pcDNA3.1-nCoV-SΔ18. HEK-293T cells were seeded one day before transfection, and the indicated plasmids were then delivered into the cells using TransIT^®^-LT1 transfection reagent (Mirus). The culture medium was refreshed at 16 h and then harvested at 48 h and 72 h post-transfection. Cell debris was removed by centrifugation at 4,000 *g* for 10 min, and the supernatant was passed through a 0.45-μm syringe filter (Pall Corporation). The pseudotyped lentivirus was aliquoted and stored at −80 °C. Transduction units (TU) of pseudotyped SARS-CoV-2 lentivirus were estimated by using a cell viability assay in response to limited dilutions of lentivirus. In brief, HEK-293T cells stably expressing the *hACE2* gene were plated on a 96-well plate 1 day before lentivirus transduction. To titer pseudotyped lentivirus, different amounts of lentivirus were added into the culture medium containing polybrene (final concentration = 8 μg/ml). Spin infection was carried out at 1,100 *g* in a 96-well plate for 30 mins at 37 °C. After incubating the cells at 37 °C for 16 h, the culture medium containing virus and polybrene was removed and replaced with fresh complete DMEM containing 2.5 μg/ml puromycin. After puromycin treatment for 48 h, the culture medium was removed and cell viability was determined using 10% AlarmaBlue reagents according to the manufacturer’s instructions. The survival rate of uninfected cells (without puromycin treatment) was set as 100%. Virus titer (TU) was determined by plotting surviving cells against diluted viral dose.

### Primary neuronal culture, pseudovirus challenge and immunostaining

Mouse neuronal cultures were prepared from dorsal cerebral cortex and hippocampal regions of WT and transgenic mice at embryonic day 16-17, as described previously [18, 19]. Embryos of both sexes were used. At days in vitro 10 (DIV 10), neurons were challenged with pseudotyped SARS-CoV-2 lentivirus (reporter: GFP) at a multiplicity of infection (MOI) of 0.01 and kept at 37 °C for 8 days. Neuronal cultures were then fixed for immunofluorescence staining as described previously [18, 19] using the following primary antibodies: rabbit anti-ACE2 (Abcam, ab108209); mouse anti-HA (Abcam, ab130275); rabbit anti-HA (Cell Signaling, 3742); mouse anti-MAP2 (Sigma, M4403); mouse anti-GFAP (Millipore, MAB3402); and rabbit anti-GFP (Invitrogen, A6455). The fluorescent images were captured at room temperature with a confocal microscope (LSM 700, Zeiss) equipped with a 20×/NA 0.80 (Plan-Apochromat) objective lens and Zen acquisition and analysis software (Zeiss). The images were processed using Photoshop (Adobe) with minimal adjustment of brightness or contrast applied to the entire images.

### Electrophysiological recording

To measure miniature excitatory postsynaptic currents (mEPSCs), we subjected cultured neurons (DIV 17-19) expressing HA tag or D614 Spike protein to whole-cell voltage-clamp recording at room temperature (23 ± 2 °C). Neurons growing on coverslips were transferred to a submerged chamber and continuously perfused with bath solution (pH 7.3) containing (in mM): 136.5 NaCl, 5 HEPES, 5.56 glucose, 5.4 KCl, 1.8 CaCl_2_, 0.53 MgCl_2_. HA tag-or Spike protein-expressing neurons were visually identified via GFP expression under an infrared differential interference contrast (IR-DIC) microscope (SliceScope, Scientifica) coupled with an OptoLED system (Cairn Research Ltd) and connected to a CCD camera (IR-1000, DAGE-MTI). Whole-cell recordings were performed with patch pipettes (4-8 MΩ) filled with the internal solution consisting of the following (in mM): 135.25 K-gluconate, 8.75 KCl, 0.2 EGTA, 4 MgATP, 10 HEPES, 7 Na2-phosphocreatine, 0.5 Na_3_GTP (pH 7.3 with KOH). Once the recording was established in the voltage-clamp configuration (Vhold = −72 mV, near the IPSC reversal potential; [Cl-]i = 8.75 mM), we applied tetrodotoxin (TTX, 1 μM) for at least 5 mins before recording mEPSCs. During mEPSC recordings, pipette capacitance and series resistance (Rs) were compensated by 70%, and Rs was continuously monitored every 10 s. Data were discarded if Rs changed by >20% during the entire 8-10-min recording period. Data were recorded using Multiclamp 700B amplifiers (Molecular Devices), filtered at 3 kHz, and sampled at 10 kHz with a Power 1401 mk II digitizer (Cambridge Electronic Design) controlled by Signal 4 software (Cambridge Electronic Design). mEPSC events were detected and analyzed by setting the peak threshold at three times the root-mean-square noise of a 2560 ms baseline and the event kinetics as corresponding to AMPAR–mediated currents. All recordings and analyses were conducted blind to the experiential conditions (i.e., HA tag or Spike protein expression).

### Statistical analysis

Statistical analyses were carried out in Excel or GraphPad Prism 8.0 software. Experiments were performed blind by relabeling the samples with the assistance of other laboratory members or without knowing genotype. Data are presented as mean values, with numbers of individual mice or neurons assessed also indicated. P values < 0.05 were considered significant.

## Results

### Establishment of CAG-hACE2 transgenic mice

To study the impact of SARS-CoV-2 infection on various tissues, we generated hACE2 transgenic mice under the control of the CAG promoter, a hybrid promoter comprising the cytomegalovirus enhancer fused to the chicken beta-actin promoter. Since the CAG promoter is highly active in a variety of tissues, CAG-hACE2 transgenic mice would serve as a model for monitoring the effects of SARS-CoV-2 infection on various tissues, including the brain. The hACE2 transgene was tagged with a HA cassette at the C-terminal end for detection. To ensure expression of the hACE2 transgene, a chicken insulator (HS4) was inserted at both the 5’ and 3’ ends of the entire transgene cassette (**Figure 1A**). We first generated two independent CAG-hACE2 transgenic lines, i.e., GT5-027 and GT4-008, using genomic PCR with primers corresponding to the sequences of hACE2 and the HA cassette (**Figure 1A, 1B**). Since the GT5-027 line bred faster than line GT4-008, we primarily used the former line for experiments unless specified otherwise. We applied ddPCR to determine transgene copy number in line GT5-027, which revealed that it carries two copies of the transgene in its genome (**Figure 1C**).

**Figure 1.**
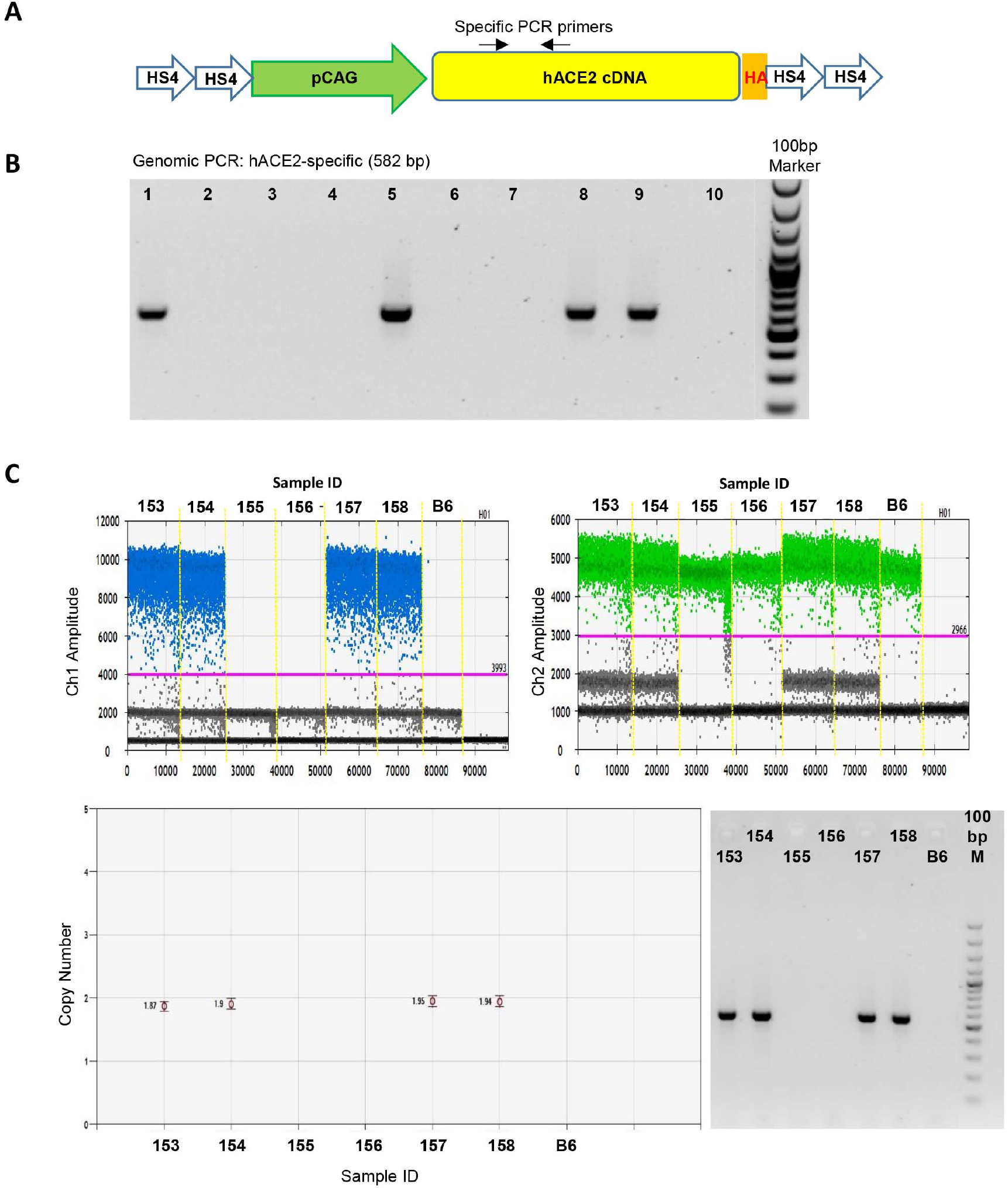
Establishment of CAG-hACE2 transgenic mice. (**A**) Transgene design for generation of CAG-hACE2 transgenic mice. Coresponding positions of *hACE2*-specific primers for genomic PCR are indicated. (**B**) Identification of CAG-hACE2 transgenic mice using genomic PCR. (**C**) Transgene copy number analysis using ddPCR. The upper panels are one-dimensional plots of droplets measured for fluorescence signal (amplitude indicated on *y*-axis) emitted from the *hACE2* transgene and *Tbpn* (reference gene). The lower panels show the copy number evaluated in QuantaSoft™ (left panel), with parallel PCR results (right panel).

### Expression of the hACE2 transgene in various organs

Expression of hACE2 transgene was examined by immunoblotting using both HA- and hACE2-specific antibodies. As expected, hACE2 proteins were detected in different organs, but with higher expression levels in the lung, kidney and brain and lower levels in the duodenum, heart and liver (**Figure 2A**). The results using hACE2 or HA antibodies were similar (**Figure 2A**). We further compared protein levels of hACE2 in male and female transgenic mice. In the six aforementioned organs, hACE2 protein levels were equivalent between female and male mice (**Figure 2B**), suggesting that both female and male transgenic mice were suitable for SARS-CoV-2 infection experiments.

**Figure 2.**
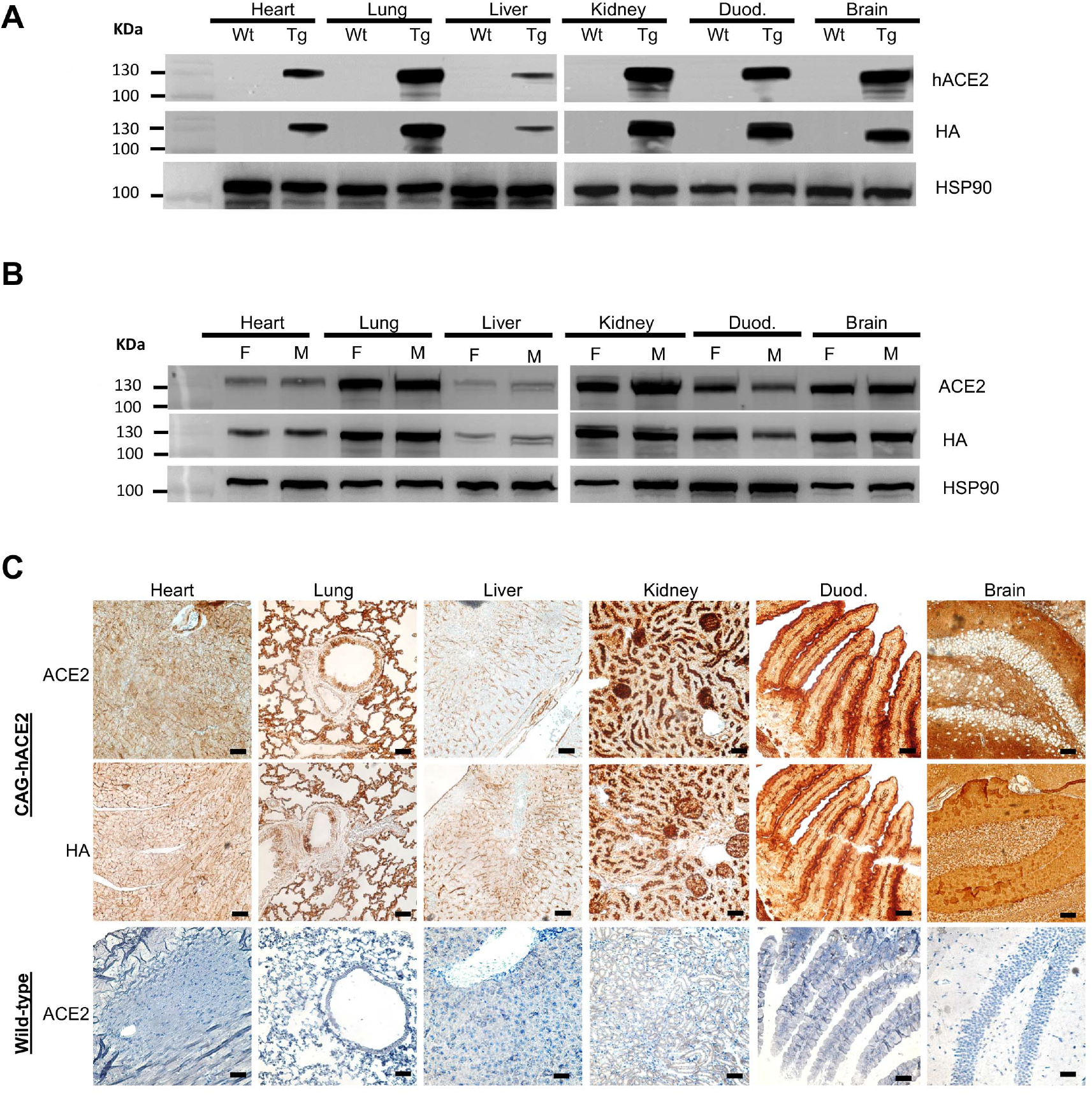
Human ACE2 protein expression in CAG-hACE2 transgenic mice. (**A**) Multiple tissue analysis of hACE2 protein expression in CAG-hACE2 transgenic mice (Tg) and WT littermates. (**B**) Multiple tissue analysis of hACE2 protein in female (F) and male (M) transgenic mice. In both (**A**) and (**B**), HA and human ACE2-specific antibodies were used to detect hACE2-HA proteins. HSP90 was used as a loading control. (**C**) Immunostaining of hACE2 protein expression in male and female tissues of WT and CAG-hACE2 transgenic mice. Scale bars, 25 mm. Duod. = duodenum.

In addition to immunoblotting, we further performed immunohistochemistry to investigate hACE2 expression at the cellular level. Patterns of HA and hACE2 immunoreactivities were generally very consistent to each other in different tissues, except for the brain (**Figure 2C**). HA antibody revealed some diffuse and non-specific background signal in the dentate gyrus (**Figure 2C**). These immunoreactivities were specific for CAG-hACE2 transgenic mice, because there was no clear hACE2 antibody signal in WT littermates (**Figure 2C**).

Thus, we have successfully generated a genetically modified mouse model that expresses hACE2 in various organs, including lung, brain, kidney, duodenum, heart and liver.

### CAG-hACE2 transgenic mice exhibit high susceptibility to SARS-CoV-2 infection

We challenged our CAG-hACE2 transgenic mice with SARS-CoV-2 via intranasal infection and then monitored changes in body weight and mouse survival (**Figure 3A**). First, we combined both the male and female transgenic mice of two mouse lines (GT5-027 and GT4-008) for comparison with their WT littermates. When challenged with 5×10^5^ PFU of SARS-CoV-2, there was no change in body weight of WT at the end of the experimental period, whereas there was a marked reduction in body weight of CAG-hACE2 mice. By day 3 post-infection (DPI 3), transgenic mice had significantly lost body weight relative to WT littermates (**Figure 3B,** unpaired t-test *p* = 0.0004), and body weight decline continued thereafter and was accompanied by a decrease in mobility. Six out of eight transgenic mice died on DPI 5 (**Figure 3B,** shown in red). Two female mice of the GT4-008 line were also challenged with the same titer of SARS-CoV-2 and showed a similar weight loss pattern (**Figure 3B**, open circles). These results suggest that our transgenic mice are susceptible to SARS-CoV-2 infection.

**Figure 3.**
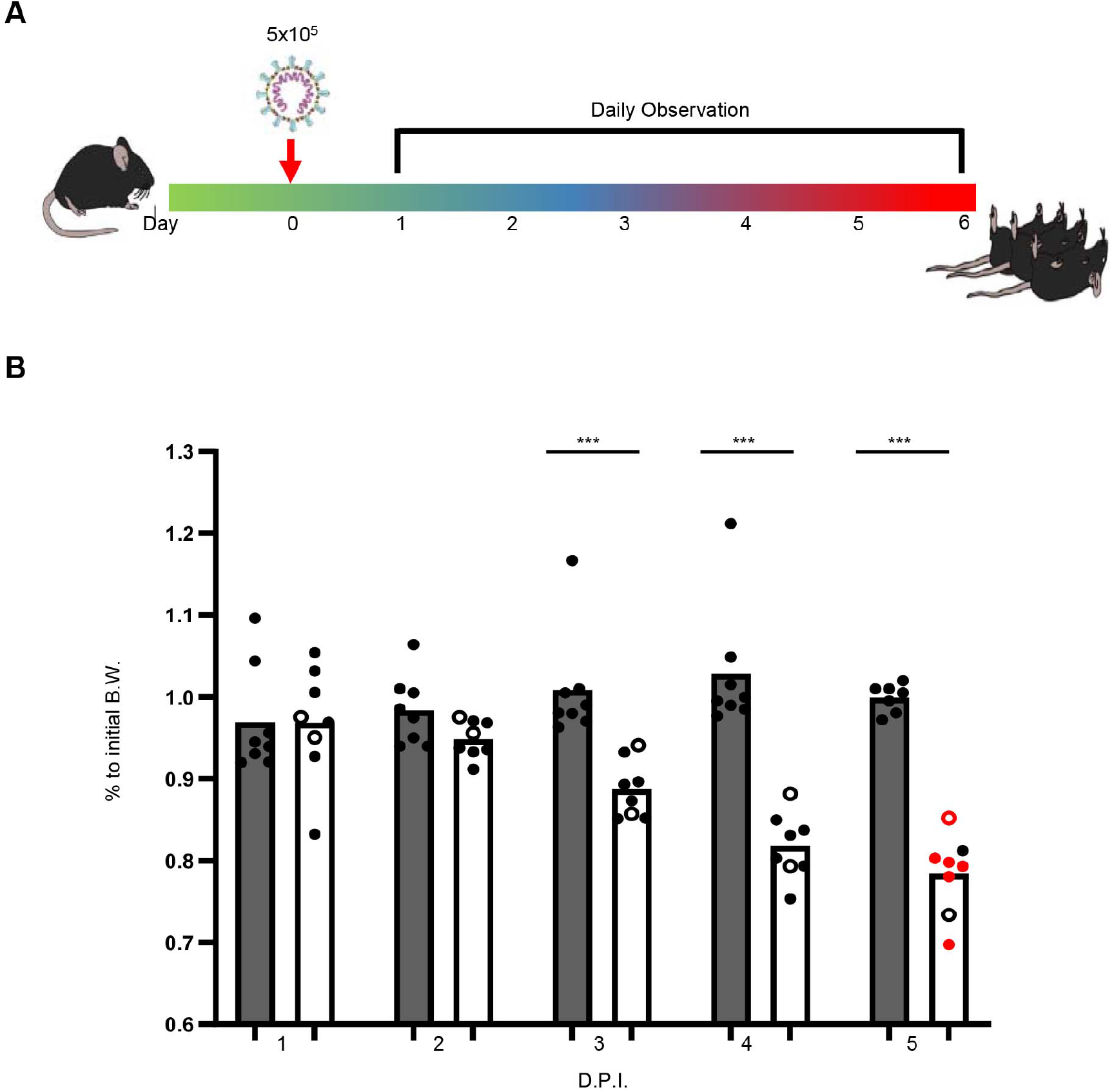
Susceptibility of CAG-hACE2 transgenic mice to SARS-CoV-2 infection. (**A**) The experimental scheme for SARS-CoV-2 infection of CAG-hACE2 transgenic mice. (**B**) Body weight changes of CAG-hACE2 transgenic mice upon SARS-CoV-2 infection at 5×10^5^ PFU. The line GT5-027 and TGT4-008 line are represented by closed and open circles, respectively. Dead mice were shown in red. Unpaired t-test: *, *p*<0.05; **, *p*<0.01; ***, p<0.005. D.P.I., days post-infection.

We reduced viral titers to determine the median lethal dose (LD_50_) of SARS-CoV-2 infection in our transgenic mice. Four different titers, i.e., 1×10^5^, 1×10^4^, 1×10^3^ and 1×10^2^, were examined (**Figure 4A**). For each titer, a total of eight transgenic mice (five males and three females of the GT5-027 line) were used. We observed that all four of these titers caused body weight loss in our transgenic mice, starting at day 3 or 4 (**Figure 4B**). Mice started to die at day 5 in the group infected with 1×10^5^, 1×10^4^ or 1×10^3^ PFU of virus, resulting in reduced mouse numbers in each experimental group (**Figure 4B**). All mice had died by DPI 9, except for one female infected with 1×10^2^ PFU (**Figure 4C, 4D**). This surviving female mouse actually fully recovered from SARS-CoV-2 infection and displayed increased body weight and normal appearance for at least two weeks after infection. Since the titer of 1×10^2^ PFU resulted in one of eight mice surviving, we estimated the LD_50_ of SARS-CoV-2 in our transgenic mice to be half the 1×10^2^ titer, i.e., 50 PFU. A total of 32 mice were used in this set of experiments, yet only one mouse survived. Thus, overall mortality (including the group of mice infected with just 100 PFU) was ~96%. Accordingly, we assert that our transgenic mice are highly susceptible to SARS-CoV-2 infection and may serve as an appropriate model for severe COVID-19.

**Figure 4.**
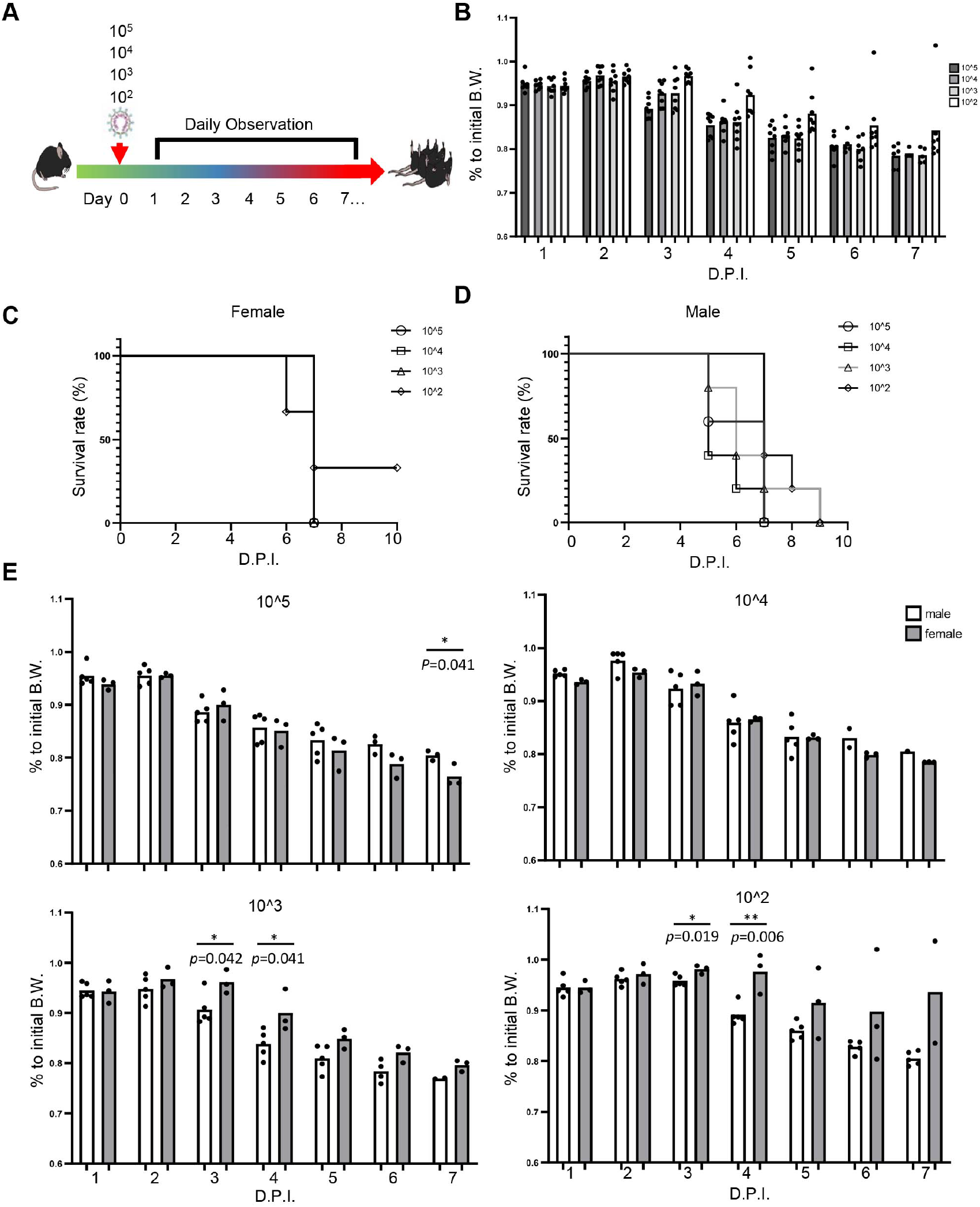
Differential responses of male and female CAG-hACE2 transgenic mice to different titers of SARS-CoV-2. **(A)** The experimental scheme for SARS-CoV-2 infection at 10^5^, 10^4^, 10^3^ and 10^2^ PFU in CAG-hACE2 transgenic mice. **(B)** Changes in body weight of CAG-hACE2 transgenic mice after infection with 10^5^, 10^4^, 10^3^ or 10^2^ PFU SARS-CoV-2. **(C), (D)** Survival curve of (C) male and (D) female CAG-hACE2 transgenic mice upon infection with 10^5^, 10^4^, 10^3^ and 10^2^ PFU SARS-CoV-2. (**E**) Changes in body weight of CAG-hACE2 transgenic mice challenged with different titers of SARS-CoV-2. Five male and three female mice were assessed for each titer. Bars represent mean value and dots show data for individual mice. Unpaired t-test: *, *p* < 0.05; **; *p* < 0.01.

### Sex-biased responses among CAG-hACE2 transgenic mice to SARS-CoV-2 infection

When we analyzed the LD_50_ of SARS-CoV-2 in our transgenic mice, we noticed that female and male transgenic mice appeared to respond differentially to viral infection. To investigate this point further, we generated survival curves for male and female transgenic mice separately (**Figure 4D**). Among the 25 male mice infected with different virus titers, six died at DPI 5. A further 3 male and 8 male mice died on days 6 and 7 post-infection, respectively. Thus, survival rates for days 5, 6 and 7 were 56%, 44% and 12%, respectively. Eventually, all infected male mice had died by DPI 9 (**Figure 4D**). In contrast, among the total of 12 experimental female mice, only one had died by day 6. However, a further 10 female mice (83% of infected mice) suddenly died on day 7, regardless of viral titer, and only one female survived SARS-CoV-2 challenge (**Figure 4D**).

We further analyzed changes in body weight of male and female mice. We found that for higher titers, i.e., 1×10^5^ and 1×10^4^ PFU, changes in body weight were comparable between male and female transgenic mice, except for the titer of 1×10^5^ at DPI 7 (**Figure 4E**). For lower titers, i.e., 1×10^3^ and 1×10^2^ PFU, female transgenic mice exhibited a much less pronounced reduction in body weight compared to male mice (**Figure 4E**). These results suggest that female and male transgenic mice respond differently to SARS-CoV-2 infection.

### Susceptibility of brain cells to SARS-CoV-2 infection

In the aforementioned immunohistochemistry study, we found that HA tag antibody revealed some non-specific signal in brain sections (**Figure 2C**). To further confirm hACE2 expression in neurons and glial cells, we prepared neuronal culture using CAG-ACE2 transgenic mice for immunofluorescence staining. Although neurons represent the majority of cell types in our neuronal culture, a small population of astrocytes is also present [20]. We first used hACE2 and HA antibodies to perform dual immunostaining on our neuronal cultures at DIV 10. Compared with WT neuronal culture, which did not express the hACE2 transgene, we observed a punctate pattern of colocalized hACE2 and HA immunoreactivities in our neuronal culture from transgenic mice (**Figure 5**), supporting that the hACE2 transgene is expressed in brain cells. Based on cell morphology, we speculated that both neurons and astrocytes expressed hACE2 (**Figure 5**, middle panels = neuron; lower panels = astrocyte). To confirm these cell types, we performed dual immunostaining using a combination of hACE2 or HA antibodies with the neuronal marker MAP2 or the astrocyte marker GFAP. Dual immunostaining using these markers indeed indicated that both neurons and astrocytes of our transgenic mice are hACE2-positive (**Figure 6A, 6B**). When we compared the immunoreactivities of neurons and astrocytes, we found that astrocytes presented much higher levels of hACE2 proteins (**Figure 6A, upper**). Thus, even though CAG is a ubiquitous promoter, the transgene expression driven by it varies. Nevertheless, these immunostaining experiments confirm expression of the hACE2 transgene in both neurons and glial cells.

**Figure 5.**
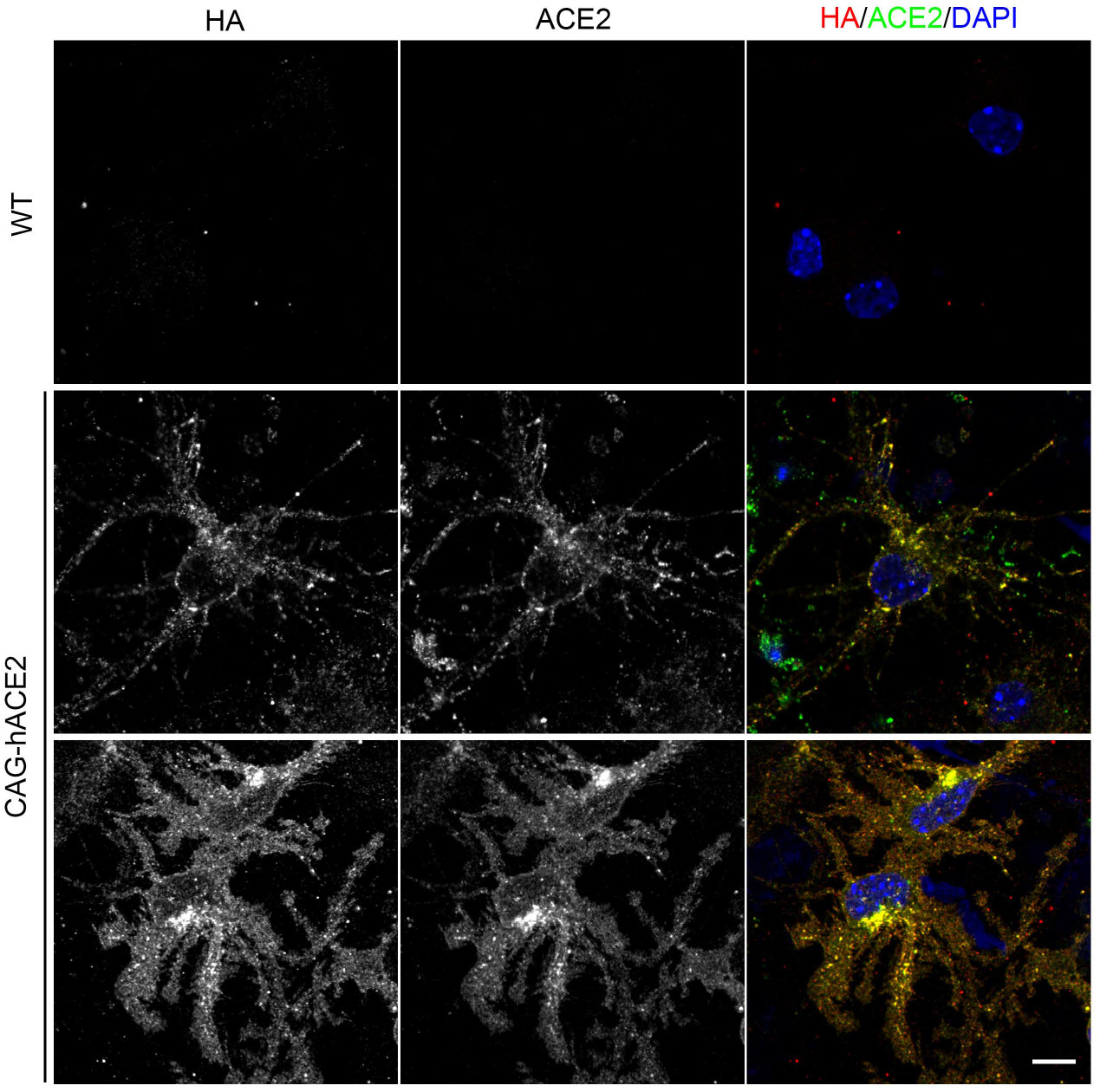
The hACE2 transgene is expressed in primary neuronal culture. Neuronal cultures were prepared using the dorsocaudal cortex and hippocampus of hACE2 transgenic mice and WT littermates. At DIV 10, WT and hACE2 transgenic neuronal cultures were subjected to immunofluorescent staining with HA and hACE2 antibodies to investigate hACE2 expression. Colocalization of HA and hACE2 immunoreactivities supports the specificity of immunostaining. Scale bar, 10 μm.

**Figure 6.**
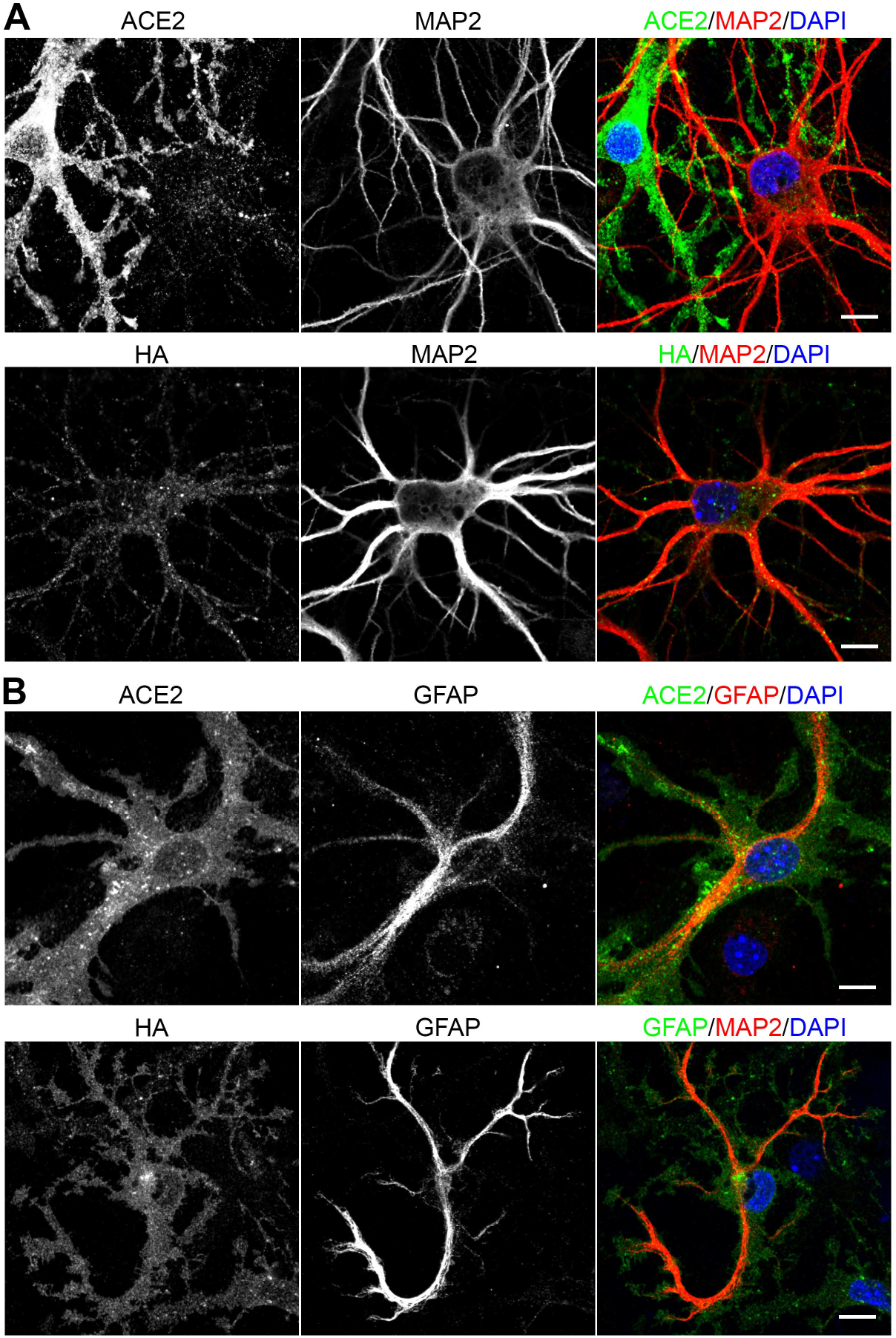
The hACE2 transgene is expressed in both neurons and astrocytes. Similar to Figure 5, neuronal cultures at DIV 10 were subjected to immunostaining using hACE2 (**A** and **B** upper panel) or HA (**A** and **B** lower panel) antibody, combined with neuronal marker MAP2 (**A**) or astrocyte marker GFAP (**B**) antibody. Scale bar, 10 μm.

We then subjected our neuronal culture to pseudovirus infection to evaluate its susceptibility to infection. To do that, we added genetically-modified lentivirus that expresses SARS-CoV-2 Spike protein co-expressed with a GFP marker into neuronal culture at DIV 10. Eight days later, cultures were subjected to immunostaining using GFP and MAP2 or GFAP antibodies (**Figure 7**). GFP expression was readily observed in both neurons and astrocytes (**Figure 7A, 7B**), indicating that our CAG-hACE2 transgenic mice can also serve as a model for investigating SARS-CoV-2 infection of the brain.

**Figure 7.**
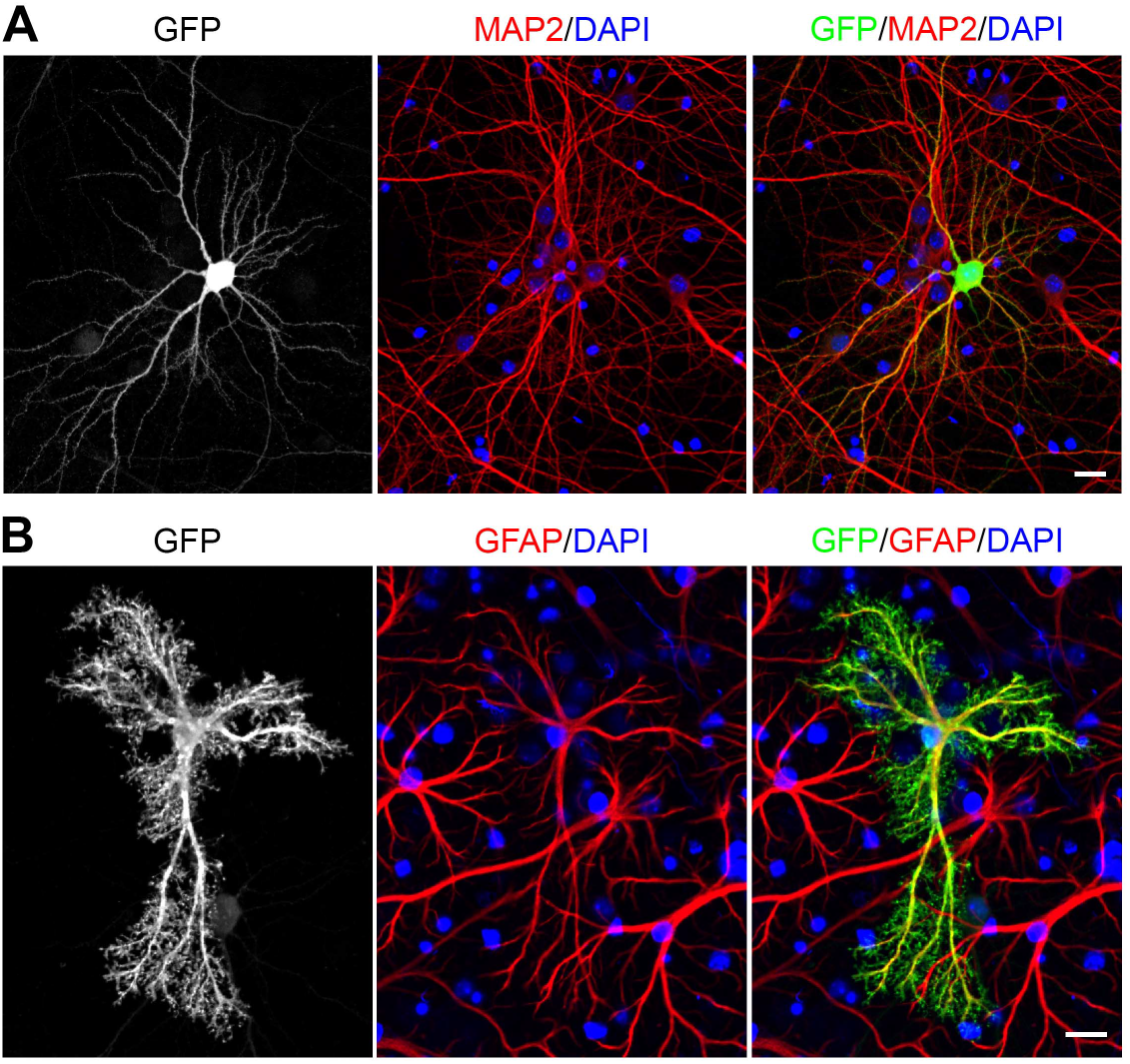
Infection of a pseudovirus expressing SARS-CoV-2 Spike protein into hACE2 transgene-expressing neurons and astrocytes. At DIV 10, neuronal cultures of hACE2 transgenic mice were challenged with SARS-CoV-2 Spike pseudovirus carrying GFP as a reporter. Cells were fixed eight days later and immunostained using GFP antibody combined with MAP2 (**A**) or GFAP (**B**) antibodies. MAP2 and GFAP are neuronal and astrocyte markers, respectively. Scale bar, 20 μm.

### SARS-CoV-2 Spike protein alters synaptic activity

Next, we investigated if SARS-CoV-2 influences neuronal activity. Due to biosafety regulations, it is technically difficult to record the electrophysiological activity of SARS-CoV-2-infected neurons. Since expression of SARS-CoV-2 Spike protein alone is sufficient to alter the morphology and density of dendritic spines, including induction of greater spine density, longer spines and narrower spine heads [15], we recorded the mEPSCs of cultured neurons transfected with SARS-CoV-2 Spike protein. GFP was co-transfected with SARS-CoV-2 Spike protein or vector control into cultured neurons to label transfected cells. Compared with vector control, we found that overexpression of SARS-CoV-2 Spike protein increased mEPSC amplitude, which was reflected in both the average of mEPSC amplitude of individual neurons and the cumulative probability curve of individual peaks (**Figure 8**). However, mEPSC frequency was not altered by this treatment (**Figure 8**). These results indicate that expression of SARS-CoV-2 Spike protein influences the synaptic activity of neurons, which is likely relevant to the neurological symptoms displayed by COVID-19 patients.

**Figure 8.**
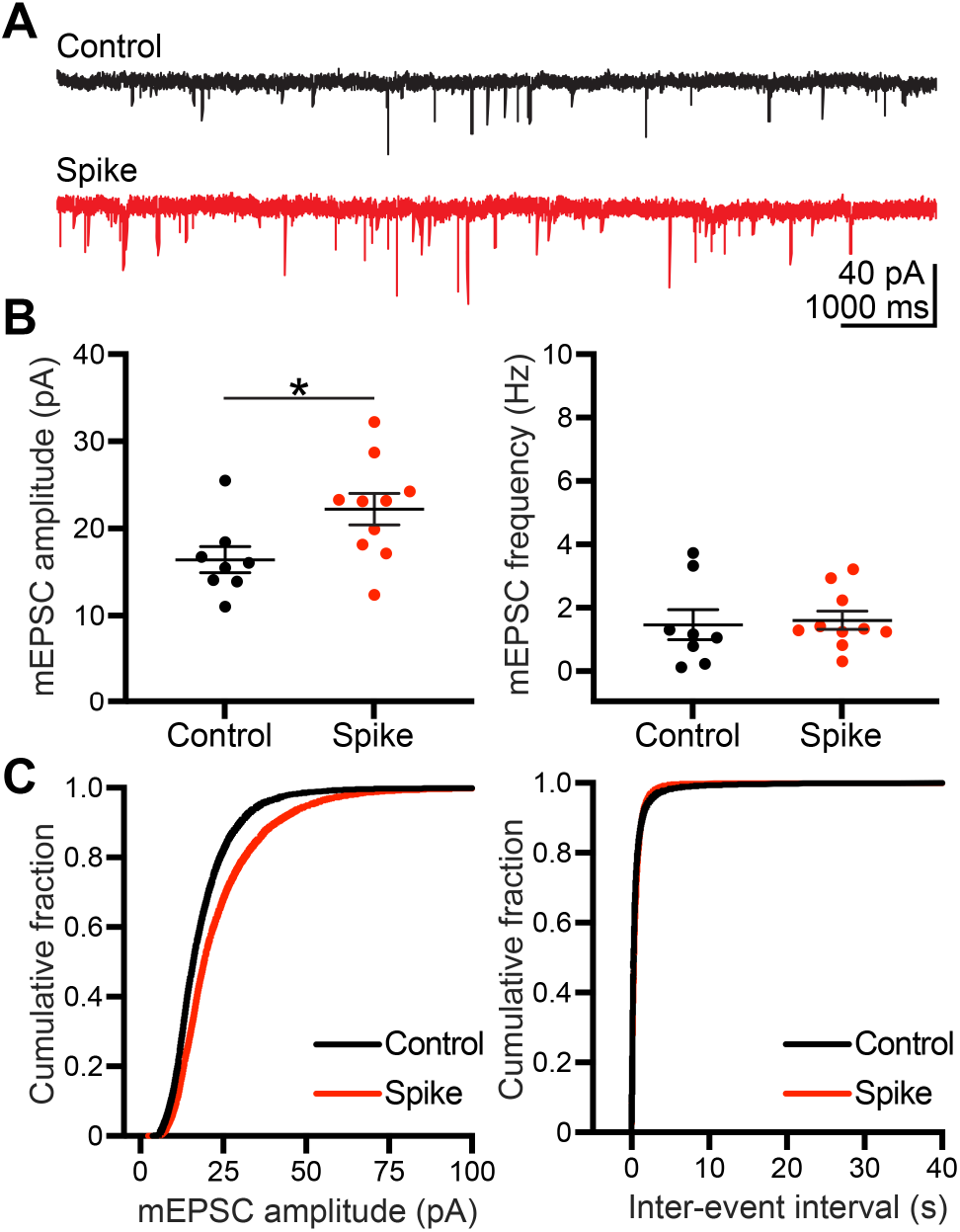
Alteration of mEPSCs by SARS-CoV-2 Spike protein. **(A)** Representative traces illustrating mEPSCs of cultured neurons transfected with vector control (black) or SARS-CoV-2 Spike protein (red). (**B**) Summary plots of average mEPSC amplitude (left) and average mEPSC frequency (right) in control (n = 8) and Spike groups (n = 10). Each dot indicates the result of an individual recording. Mean ± SEM are also shown. Mann-Whitney U test; * *p* < 0.05. (**C**) Cumulative distribution of mEPSC amplitude (left) and inter-event intervals (right) recorded in control and Spike groups.

## Discussion

Our hACE2 transgene was driven by a ubiquitous promoter, CAG, and was flanked with two copies of the HS4 insulators to limit transgene silencing by positional effects. This design resulted in strong expression of hACE2 in different organs of our transgenic mice. Thus, virus propagation to other organs upon intranasal infection is expected to occur in this mouse model. However, we noticed that although the CAG promoter is ubiquitous, hACE2 expression levels vary across different tissues. In neuronal culture, we also observed that astrocytes and neurons express very different levels of hACE2 proteins. It is possible that hACE2 protein is subject to posttranslational regulation. Nevertheless, the higher expression levels of hACE2 in the astrocytes of our transgenic mice may indicate that astrocytes display higher viral susceptibility, mimicking the tropism of SARS-CoV-2 reported for human cortical astrocytes in organoids [21]. Our CAG-hACE2 transgenic mice also express hACE2 in diverse tissues, making them a useful model for investigating systematic responses to SARS-CoV-2 infection.

We have also demonstrated that our CAG-hACE2 transgenic mouse line is a very sensitive model for SARS-CoV-2 infection, as it responded to very low titers of SARS-CoV-2. The LD_50_ of SARS-CoV-2 in our hACE2 transgenic mice was lower than 10^2^ PFU and we estimated it to be ~50 PFU, which is the lowest LD_50_ yet reported for SARS-CoV-2. Typically, 2.5-10 x 10^4^ PFU of SARS-CoV-2 are used for infection experiments using mouse models [6, 12, 13]. For our CAG-hACE2 transgenic mice, a titer of 100 PFU was sufficient to cause 100% lethality in male mice. Thus, relative to other currently available models, our hACE2 transgenic mice are the most sensitive model for COVID-19 infection. Interestingly, female hACE2 transgenic mice are relatively resistant to low-dose SARS-CoV-2 infection, displaying a lower degree of body weight loss compared to males and one female infected with 100 PFU even survived at least longer than 2 weeks after infection. This difference is unlikely due to differential expression levels of hACE2 proteins in male and female mice because immunoblotting revealed comparable hACE2 protein levels among male and female mice for various organs. Thus, our transgenic mice may also serve as a model to study severe COVID-19 and sex-biased responses to SARS-CoV-2 infection. Consequently, it should prove valuable in exploring therapeutic agents and for vaccine development.

Our previous report showed that overexpression of SARS-CoV-2 Spike protein in cultured neurons alters dendritic spine density and morphology [16]. Since dendritic spines are mainly supported by F-actin cytoskeleton, this alteration of dendritic spines suggests that SARS-CoV-2 can influence F-actin. Consistent with this observation, informational spectrum analysis has revealed that actin is a possible host factor for cell entry and pathogenesis of SARS-CoV-2 [22]. Confocal analysis has further indicated that SARS-CoV-2 Spike protein colocalizes with F-actin along filopodia, another subcellular structure supported by F-actin in non-neuronal cells [16]. The presence of SARS-CoV-2 virions along filopodia may facilitate virus spread [23]. Here, we further show that the mEPSCs of SARS-CoV-2 Spike protein-expressing neurons differs from those of control neurons, further implying that SARS-CoV-2 infection alters neuronal activity, which may account at least partially for the neurological symptoms displayed by COVID-19 patients. Apart from neurons, the astrocytes of our transgenic mice were also readily infected by SARS-CoV-2. Thus, our transgenic mice provide an appropriate model for investigating the impact of SARS-CoV-2 infection on various organs and tissues, as well as the neuropsychiatric symptoms observed in COVID-19 patients [24, 25].

## Acknowledgement

We thank the COVID-19 Team of Academia Sinica, and the Animal Facility and Genomic Core Facility of the Institute of Molecular Biology, Academia Sinica for technical support. Dr. John O’Brien conducted English editing. Members of Dr. Yi-Ping Hsueh’s laboratory relabeled samples for blind experiments. This work was supported by the Core Facilities and Innovative Instrument Project, Academia Sinica (AS-CFII-108-104 to C.-Y.T.) and the Institute of Molecular Biology, Academia Sinica to Y.-P.H..

